## Supplementary figures and tables for "ultraID: a compact and efficient enzyme for proximity-dependent biotinylation in living cells"

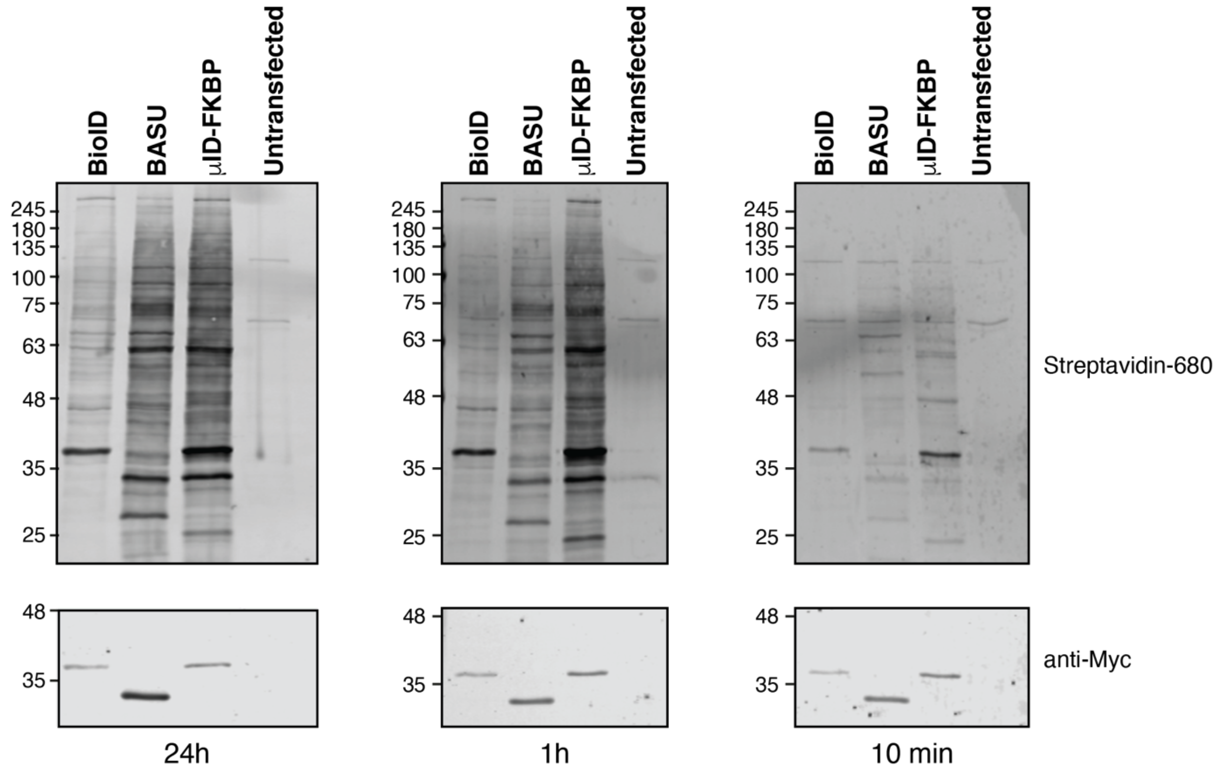


**Supplementary Figure 1: microID is an enhanced activity PDB enzyme.**
Blots of lysates of HeLa cells transiently transfected with the indicated constructs and incubated with 50 µM biotin for 24 h (left), 1 h (middle) or 10 min (right). Biotinylation was analyzed using IRDye680-labeled streptavidin and expression levels of the fusion proteins with antibodies against the Myc tag as indicated.

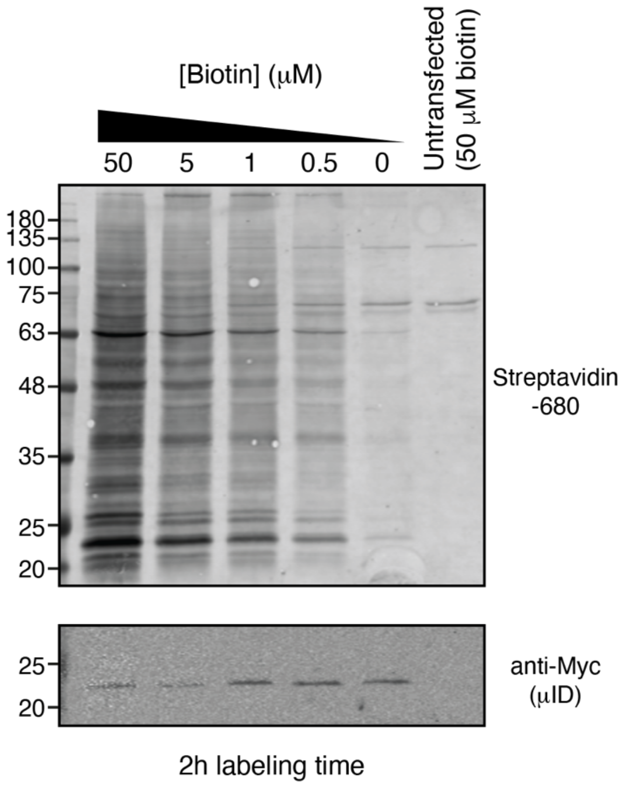


**Supplementary Figure 2: microID requires lower concentrations of biotin.**
Blots of lysates of HeLa cells transiently expressing Myc-tagged microID and incubated with the indicated concentrations of biotin for 2 h. Biotinylation was analyzed using IRDye680-labeled streptavidin and expression levels of the fusion proteins with antibodies against the Myc tag as indicated.

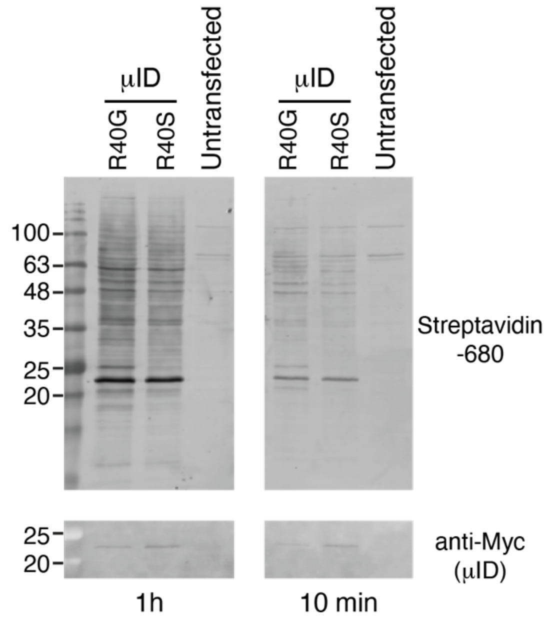


**Supplementary Figure 3: The R40S mutation does not enhance the activity of microID.**

Blots of lysates of HeLa cells transiently expressing Myc-tagged microID with either the R40G or the R40S mutations and incubated with with 50 μM biotin for 1 h (left) or 10 min (right). Biotinylation was analyzed using IRDye680-labeled streptavidin and expression levels of the fusion proteins with antibodies against the Myc tag as indicated.

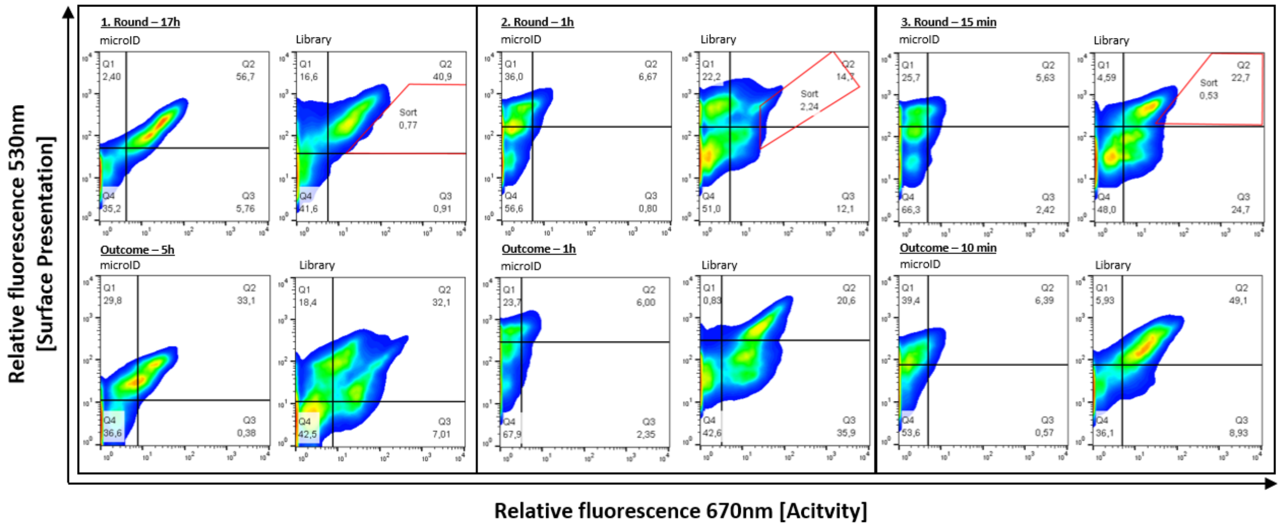


**Supplementary Figure 4: Progress of the directed evolution of microID.**
Upper row: sorting rounds with indication of the sorting cells (red gates), bottom row: outcome of the cell enrichment after sorting.

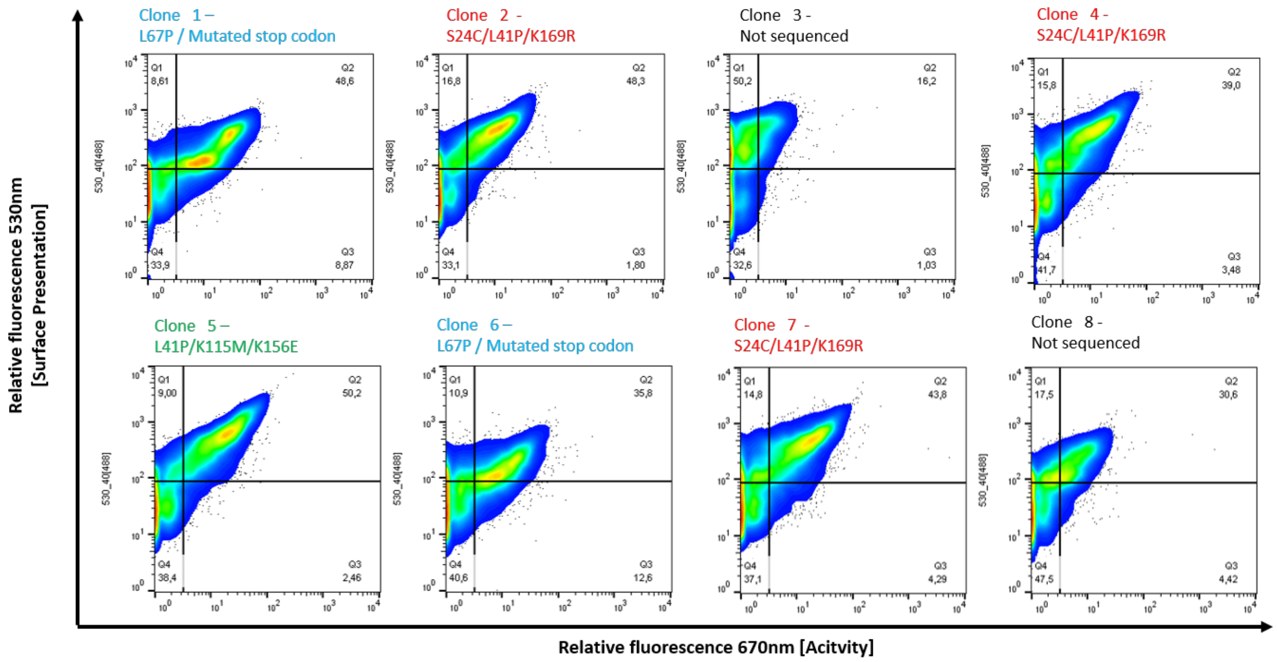


**Supplementary Figure 5: Single clone analysis from the round 3 yeast library.** The indicated clones were analyzed for surface presentation (y-axis) and biotinylation activity (x-axis). The mutations, deduced by DNA sequencing, on each clone are indicated.

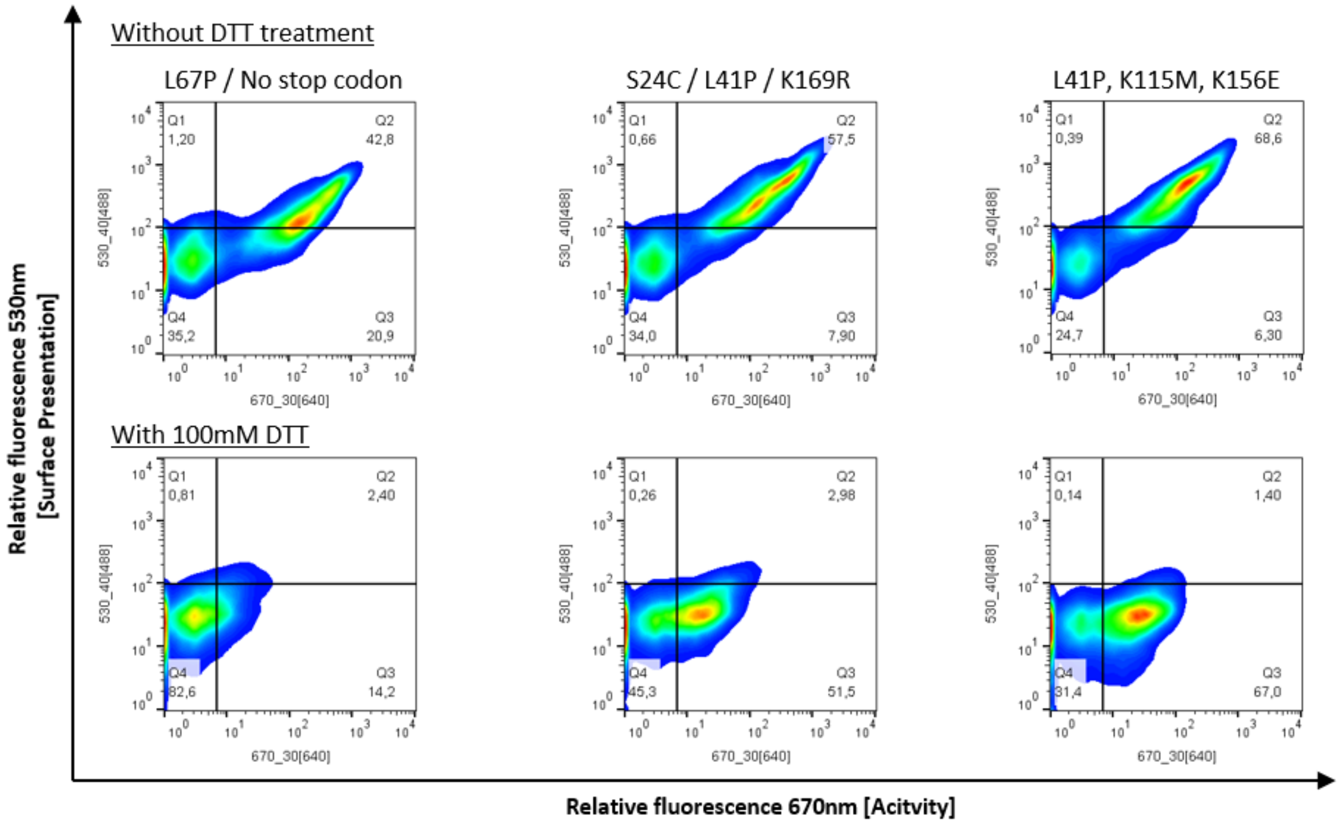


**Supplementary Figure 6: Proximity-dependent biotinylation test for the potentially improved microID variants.**
The indicated clones were analyzed for surface presentation (y-axis) and biotinylation activity (x-axis). Upper row: analysis after the cell surface biotinylation assay. Lower row: analysis after the cell surface biotinylation and release of the enzymes by reducing the disulfide bonds between Aga1p and Aga2p with DTT.

A

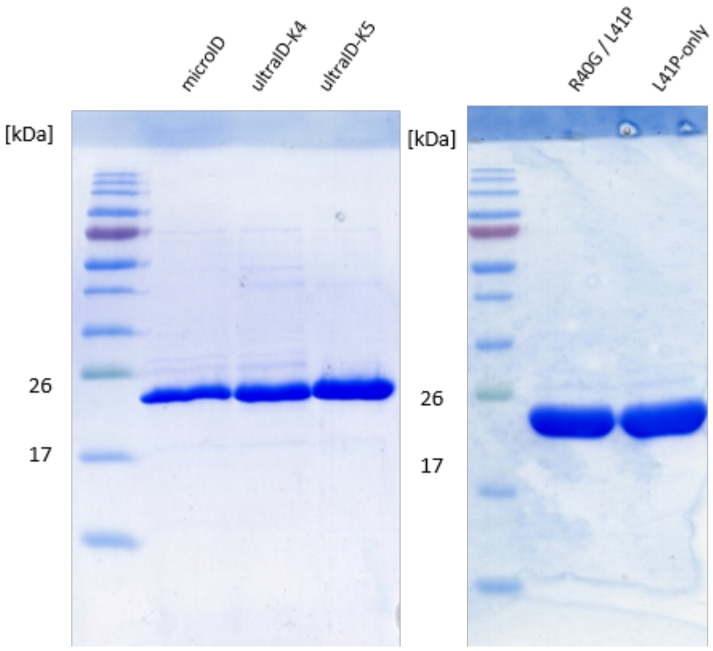


B

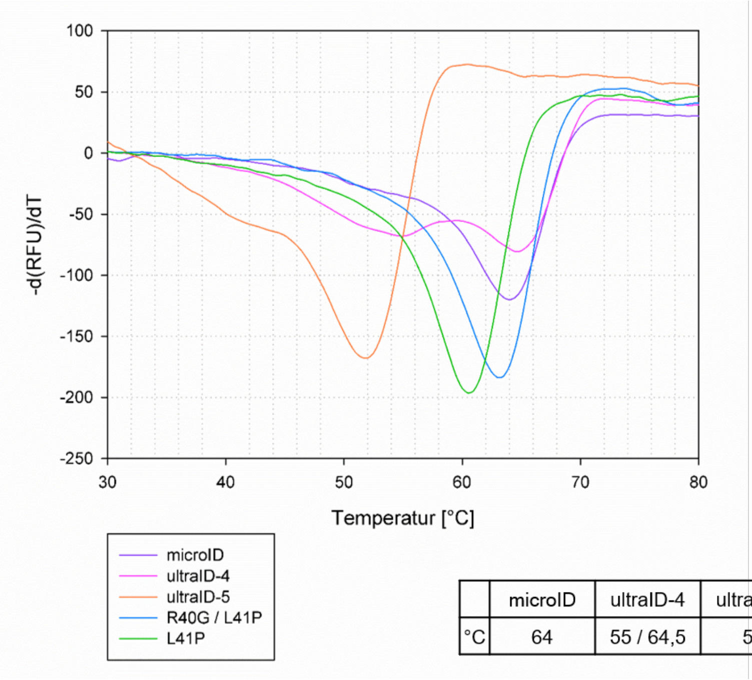


| | microID | ultraID-4 | ultraID-5 | R40G / L41P | L41P |
|---|---|---|---|---|---|
| °C | 64 | 55 / 64,5 | 52 | 63 | 60,5 |

**Supplementary Figure 7: Recombinant microID variants and their measured thermostability.**

(A) Picture of a Coomassie blue-stained acrylamide gel showing the indicated IMAC affinity purified proteins. (B) Thermostability profile of the purified proteins from (A) determined with a SYPRO orange-based thermal shift assay.

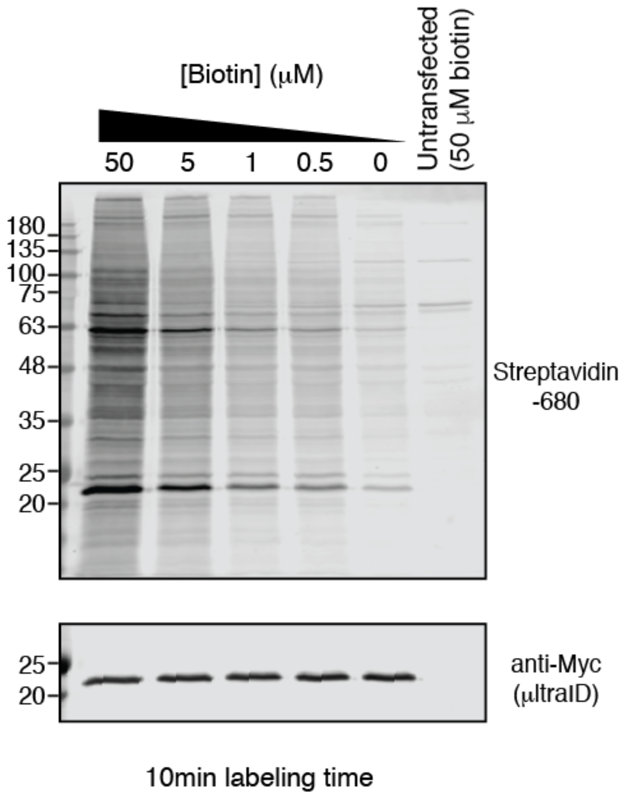


**Supplementary Figure 8: ultraID requires lower concentrations of biotin.**
Blots of lysates of HeLa cells transiently expressing Myc-tagged ultraID and incubated with the indicated concentrations of biotin for 10 min. Biotinylation was analyzed using IRDye680-labeled streptavidin and expression levels of the fusion proteins with antibodies against the Myc tag as indicated.

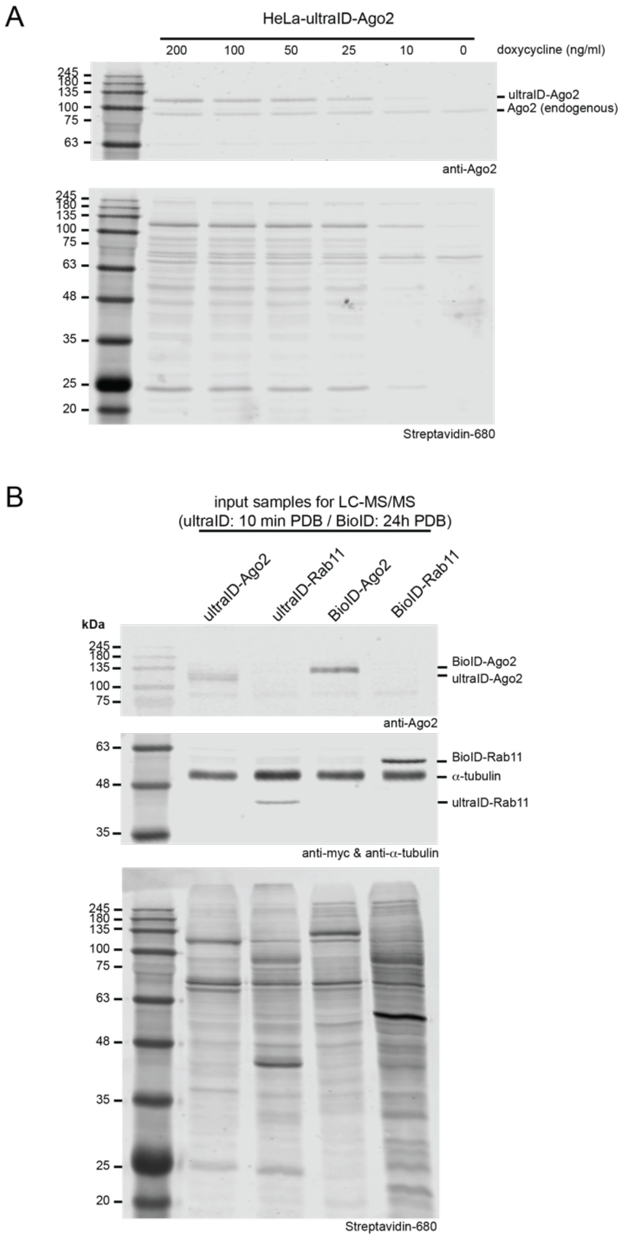


**Supplementary Figure 9: Efficient biotinylation with ultra-Ago2 at physiological expression levels.**

(A) Blots of lysates of HeLa cells stably expressing Myc-tagged ultraID-Ago2 after induction with the indicated concentrations of doxycycline. PDB was induced with biotin for 10 min. (B) Blots of lysates of HeLa cells stably expressing the indicated fusion proteins and used as a starting material for the LC-MS/MS analysis. Biotinylation was analyzed using IRDye680-labeled streptavidin and expression levels of the fusion proteins with antibodies against the Myc tag or Ago2 as indicated. Detection of $\alpha$-tubulin serves as a loading control on the middle panel. Upper and lower panels belong to the same membrane analyzed with two fluorophores, the endogenous biotinylated protein running between the 63 and 75 kDa markers serves as an internal loading control.

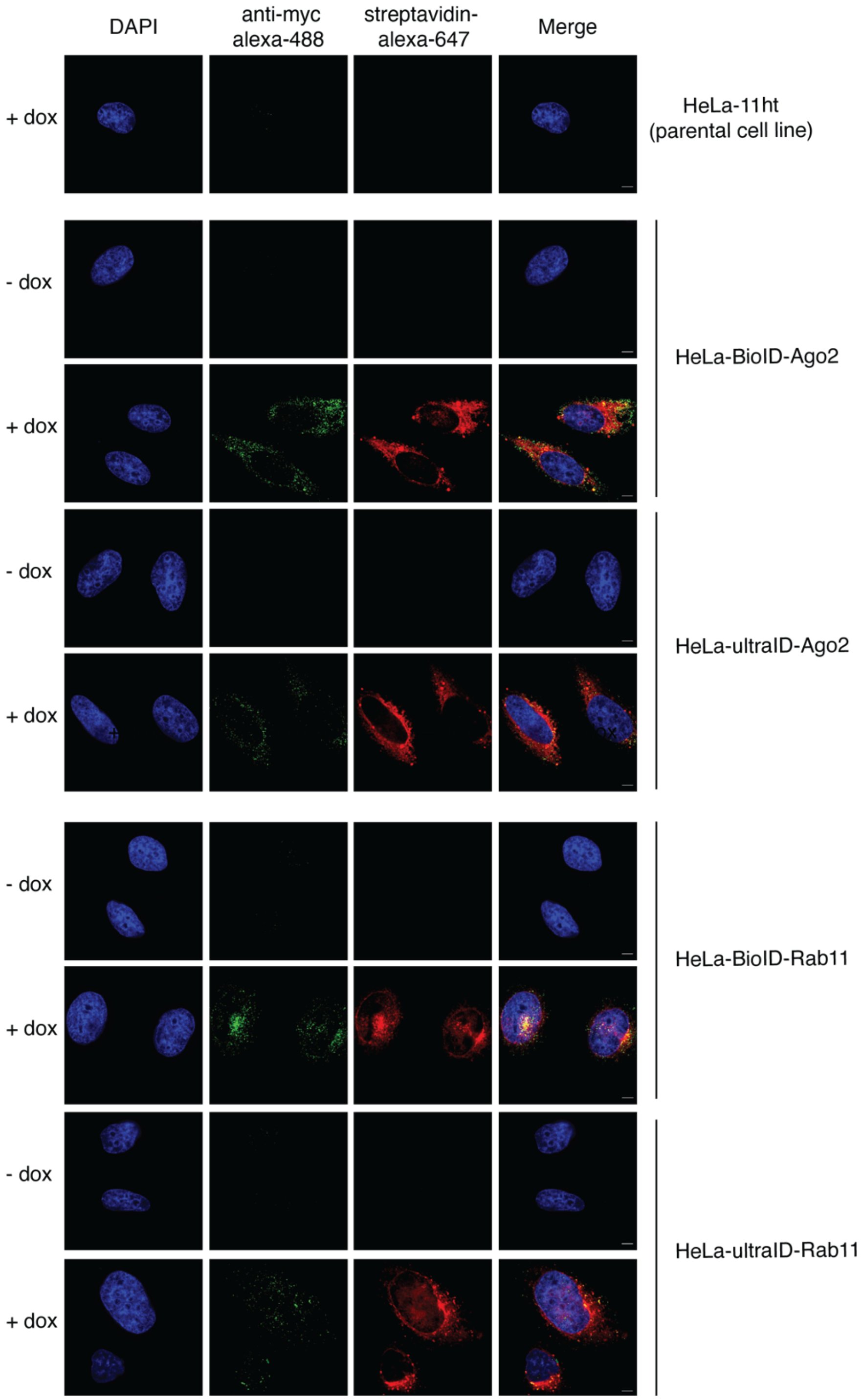


**Supplementary Figure 10: Localization of the ultraID-Ago2/Rab11 and BioID-Ago2/Rab11 fusion proteins.**
Immunofluorescence of the indicated expressed fusion proteins by Myc-tag detection as well as the detection of biotinylated proteins by alexa-488-coupled streptavidin. Scale bar, 5 µm. All fusion proteins are expressed in the cytosol and show their expected staining pattern.

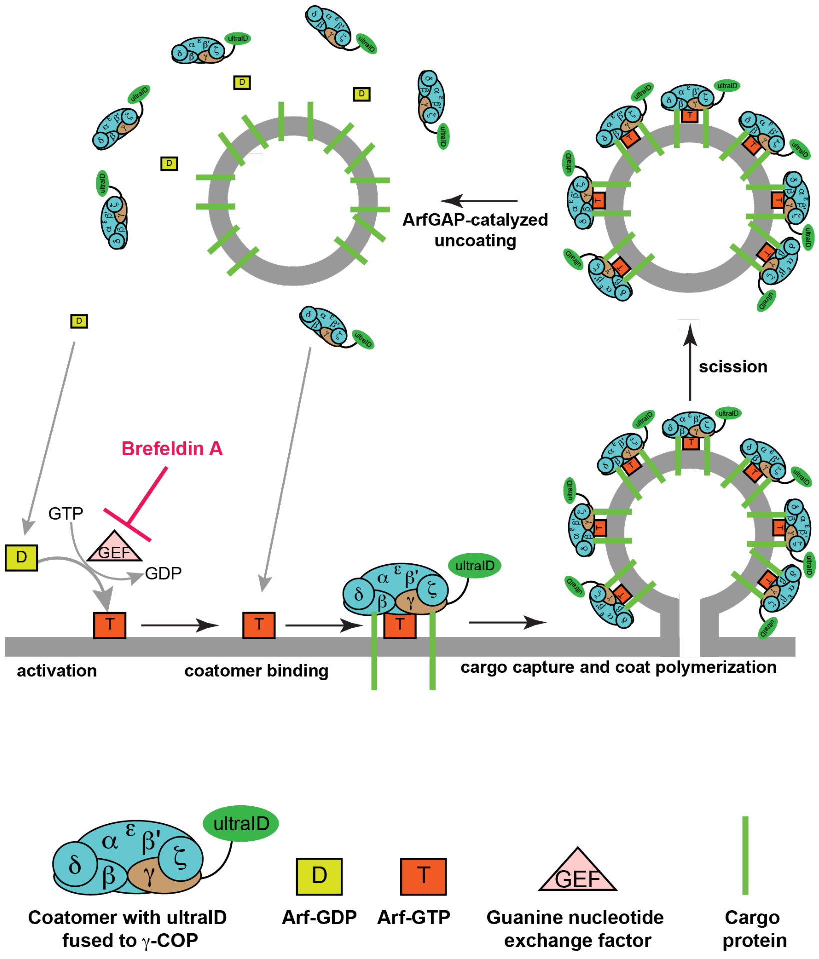


**Supplementary Figure 11: Cycle of coatomer recruitment to and release from Golgi membrane.**
Cytosolic coatomer is recruited to Golgi membranes by the small GTPase Arf after its activation through GDP to GTP exchange mediated by a specific GEF. Once at the Golgi membrane, the coatomer/Arf complex binds to transmembrane cargo proteins and polymerizes, thereby deforming the membrane until a vesicle pinches off. Soon after membrane scission, GTP-hydrolysis by Arf1 is stimulated by ArfGAPs, leading to Arf-GDP and coatomer release to the cytosol. The drug brefeldin A is a specific inhibitor of the Arf-GEF, thereby blocking the recruitment of coatomer to membranes.

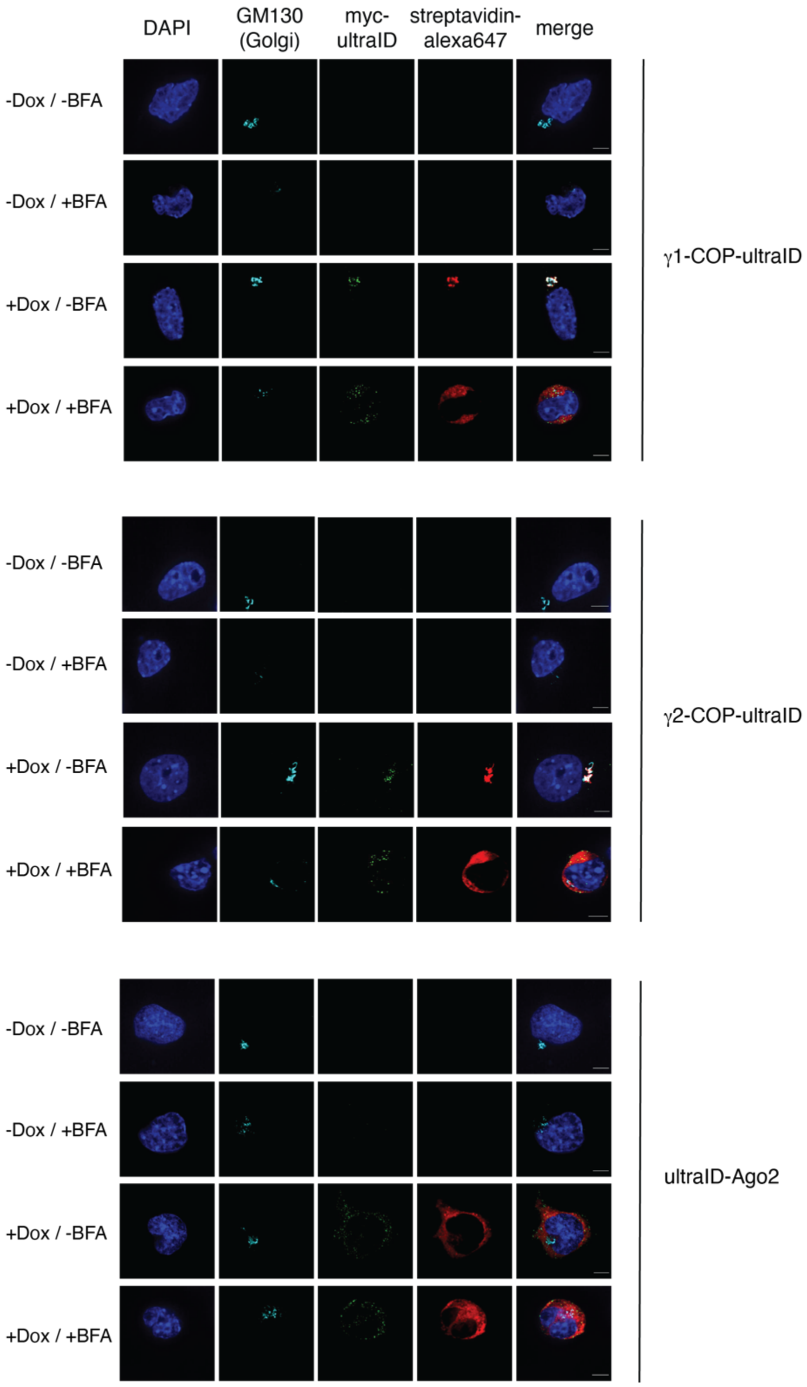


**Supplementary Figure 12: Localization of the γ1-COP-ultraID, γ2-COP-ultraID and ultraID-Ago2 fusion proteins.**
Immunofluorescence of the indicated expressed fusion proteins by Myc-tag detection and of the Golgi marker GM130. Biotinylated proteins after PDB were detected with alexa-647-coupled streptavidin. Scale bar, 5 µm. The non-induced cell lines (-Dox) served as a staining specificity control. When indicated the cells had been treated with brefeldin A (+BFA).

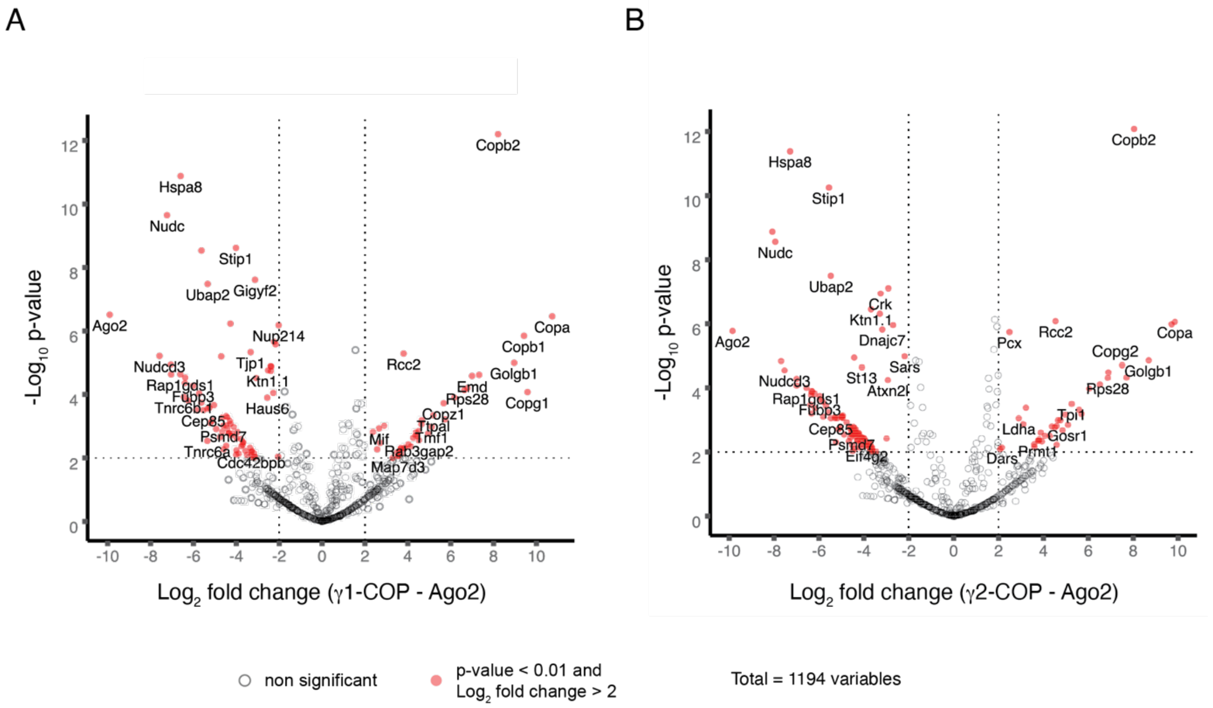


**Supplementary Figure 13: cytosolic and membrane proteins proximal to γ1-COP and γ2-COP.**

(A) Volcano plots of the proteins identified by LC-MS/MS for the γ1-COP-ultraID protein after PDB under mock conditions. The ultraID-Ago2 dataset served as a negative control. Significant hits (adjusted p-value < 0.01 and $log_2$ fold change > 2) are indicated in red. Selected hits are labeled. (B) same as (A) for the γ2-COP-ultraID protein

**Supplementary Table 1: Plasmids generated in this study**

| Plasmid | Description |
|---|---|
| pSF3-NBioID2-FKBP/CBioID2-FRB | Co-expression of NBioID2-FKBP and CBioID2-FRB in HeLa-11ht cells |
| pSF3- NBioID2-FKBP | Expression of NBioID2-FKBP in HeLa-11ht cells |
| pSF3-BioID | Expression of BioID in HeLa-11ht cells |
| pSF3-BioID2 | Expression of BioID2 in HeLa-11ht cells |
| pSF3-BASU | Expression of BASU in HeLa-11ht cells |
| pSF3-TurboID | Expression of TurboID in HeLa-11ht cells |
| pSF3-microID | Expression of microID in HeLa-11ht cells |
| pSF3-microID-L41P only | Expression of microID-L41P in HeLa-11ht cells |
| pSF3-ultraID-4 | Expression of ultraID-4 in HeLa-11ht cells |
| pSF3-ultraID-5 | Expression of ultraID-5 in HeLa-11ht cells |
| pSF3-ultraID | Expression of ultraID in HeLa-11ht cells |
| pCT-clone 1 | Yeast display expression plasmid for clone 1 of the directed evolution selection |
| pCT-clone 4 (ultraID-4) | Yeast display expression plasmid for clone 4 of the directed evolution selection |
| pCT-clone 5 (ultraID-5) | Yeast display expression plasmid for clone 5 of the directed evolution selection |
| pET-22b-microID | Expression of microID in *E. coli* |
| pET-22b-microID-L41P only | Expression of microID-L41P in *E. coli* |
| pET-22b-ultraID-4 | Expression of ultraID-4 in *E. coli* |
| pET-22b-ultraID-5 | Expression of ultraID-5 in *E. coli* |
| pET-22b-ultraID | Expression of ultraID in *E. coli* |
| pET15b-CNOT9 | Expression of CNOT9 in *E. coli* |
| pET15b-microID | Expression of microID in *E. coli* |
| pET15b-ultraID | Expression of ultraID in *E. coli* |
| indPB4-$\gamma$1-COP-ultraID | PiggyBac-based plasmid for making doxycycline-inducible stable P19 cells for the expression of $\gamma$1-COP-ultraID |
| indPB4-$\gamma$2-COP-ultraID | PiggyBac-based plasmid for making doxycycline-inducible stable P19 cells for the expression of $\gamma$2-COP-ultraID |
| indPB4-ultraID-Ago2 | PiggyBac-based plasmid for making doxycycline-inducible P19 cells for the expression of ultraID-Ago2 |
| pME2787 | *MET25Prom*, *CYC1Term*, *URA3,* 2 µm (reference: Mumberg et al., 1994) |
| pME4478 | *MET25Prom*, *CYC1Term*, *URA3,* 2 µm, *ASC1-birA** (reference: Opitz *et al*. 2017) |
| pME5086 | *MET25Prom*, *CYC1Term*, *URA3,* 2 µm, *ASC1-µID* |
| pME5087 | *MET25Prom*, *CYC1Term*, *URA3,* 2 µm, *ASC1-ultraID* |
| pSF3-ultraID-Ago2 | For making doxycycline-inducible stable HeLa 11ht cells for the expression of ultraID-Ago2 |
| pSF3-ultraID-Rab11 | For making doxycycline-inducible stable HeLa 11ht cells for the expression of ultraID-Rab11 |

**Supplementary Table 2: *S. cerevisiae* strains used in this work**

| Strain | Genotype | Reference |
|---|---|---|
| RH2817 | *MATα, ura3-52, trp1::hisG* | Valerius *et al*., 2007 |
| RH3263 | *MATα, ura3-52, trp1::hisG, leu2::hisG, Δasc1::LEU2* | Valerius *et al*., 2007 |

**Supplementary Table 3: Antibodies used in this study**

| Antibody | Host | Source | Dilution |
|---|---|---|---|
| Anti-myc (9E10) | mouse | DSHB (sc-40X) | WB, 1:1‘000 |
| Anti-FLAG (M2) | mouse | Sigma (F1804) | WB, 1:500 |
| Anti-CNOT9 | rabbit | Proteintech (22503-1 AP) | WB, 1 :3’000 |
| Anti-$\alpha$-tubulin (B-5-1-2) | mouse | Sigma (T5168) | WB, 1:10‘000 |
| Anti-Ago2 (11A9) | rat | Sigma (MABE253) | WB, 1:5‘000 |
| anti-$\gamma$1-COP (anti-$\gamma$1-app) | rabbit | Wieland lab (Heidelberg) | WB, 1:800 |
| anti-$\gamma$2-COP (anti-$\gamma$2-app) | rabbit | Wieland lab (Heidelberg) | WB, 1:800 |
| anti-Asc1p | rabbit | Valerius lab | WB, 1:1‘000 |
| anti-myc-Tag (71D10) | rabbit | Cell signalling technology (2278S) | IF, 1:400 |
| Anti-GM130 (35/GM130) | mouse | BD Transduction Laboratories (610822) | IF, 1:500 |
| anti-mouse IgG IRDye 800CW | goat | LI-COR (C40826-01) | WB, 1:15‘000 |
| anti-rat IgG DyLight800 | goat | Thermo Scientific (SA5-10024) | WB, 1:15‘000 |
| anti-mouse IgG AlexaFluor680 | goat | Thermo Scientific (A-21057) | WB, 1:10‘000 |
| anti-rabbit IgG IRDye 800CW | goat | LI-COR (926-32211) | WB, 1:15‘000 |
| anti-rabbit IgG IRDye 680CW | goat | LI-COR (926-68071) | WB, 1:15‘000 |
| anti-mouse Alexa 647 | donkey | Invitrogen (A28175) | IF, 1:1‘000 |
| anti-mouse Alexa 546 | goat | Invitrogen (A11030) | IF, 1:1‘000 |
| anti-rabbit Alexa 488 | goat | Invitrogen (A11008) | IF, 1:1‘000 |
| streptavidin-DyLight680 | - | Invitrogen (21848) | WB, 1:15‘000 |
| streptavidin-AlexaFluor647 | - | Jackson ImmunoResearch (016-600-084) | IF, 1:1‘000 |
